# Experimentally Tuned Protein-RNA Rosetta Score Function using Bayesian Optimization

**DOI:** 10.64898/2026.07.17.739244

**Authors:** Joseph S. Bailey, Nathan Phan, Søren C. Spina, Rachel B. Getman, Joel A. Paulson, Blaise R. Kimmel

## Abstract

Protein-RNA complexes drive fundamental cellular processes such as transcription and translation. Despite the prevalence and importance of protein-RNA interactions, the field lacks reliable and accessible methods to quantify the energetic favorability of these interactions. We propose an experimentally tuned protein-RNA score function that can be directly implemented into ROSETTA. Fine-tuning these score functions for predictive tasks requires repeated evaluations on a set of protein-RNA complexes, which can be computationally expensive given the number of parameters to tune. We used Bayesian Optimization to efficiently improve the energetic agreement between ROSETTA and experimentation. We observe significant interactions for specific RNA subclasses, serving as further confirmation of the physical validity of the score function. Beyond protein-RNA interaction prediction, we establish a framework to efficiently fine-tune ROSETTA score functions for any protein-class interaction using Bayesian Optimization.

**TOC FIGURE:** 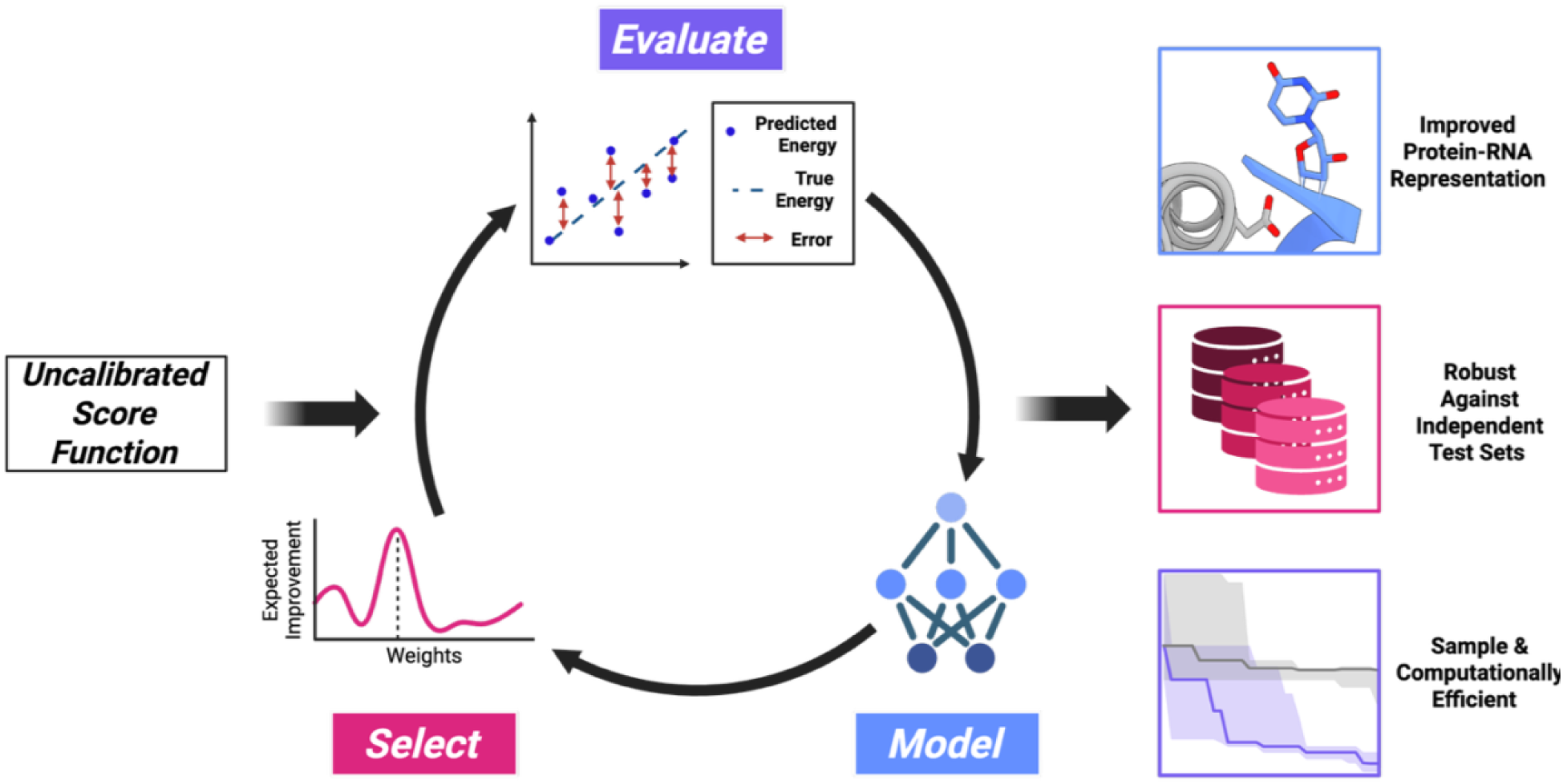

## Introduction

Protein-RNA interactions play key roles in transcription, translation, and RNA processing. Accurately measuring these interactions is the first step toward modeling and designing new functional RNA-binding proteins^1–3^. Experimental techniques provide direct measurements of protein-RNA complex binding effects; however, conducting experiments with these methods can be resource- and time-intensive^4^. This challenge is amplified by the high flexibility of RNA that makes binding patterns difficult to characterize. Hence, there is a need for computational algorithms that can efficiently score the diverse range of protein-RNA interactions consistent with experimental measurements. When used for protein-RNA interactions, existing score functions fall short of achieving high-resolution accuracy and fail to capture the key energetic factors that govern RNA recognition at the binding interface^5^, as current score functions were developed using protein or small-molecule benchmarks. In turn, it is intrinsically assumed that the energetic patterns captured in those models will behave the same way when applied to protein-RNA interactions.

The physicochemical properties of protein-RNA complexes make protein-protein score functions a poor fit for prediction, often resulting in large deviations from experimentally measured binding data^6^. For example, protein-RNA interfaces often retain water molecules, which lead to residue-length and complex-dependent behaviors that protein-trained scoring terms do not capture well. RNA backbones carry substantial charges, which lead to long-range electrostatic effects, directional, base-specific hydrogen bonds, and unique interfacial constraints caused by frequent π-stacking interactions. To overcome these challenges, recent efforts to improve binding predictions have shifted toward machine learning (ML) approaches^7,8^. These models have shown promise but offer limited insight into which underlying forces are misbalanced or how their learned corrections generalize across different protein-RNA classes^9^. Additionally, they are often difficult to apply because they rely on large decoy libraries and many iterations, resulting in high computational cost.

ROSETTA is a computational suite for modeling, predicting, and designing proteins^10,11^. It ranks biomolecular structures using tunable score functions that include physics-based terms capturing intra- and intermolecular interactions, such as van der Waals and hydrogen bonding. Fine-tuning ROSETTA score function weights is done heuristically using knowledge of interaction behavior, for example, the relative importance of hydrogen bonding. This approach, while intuitive, may fail to discover underlying dependencies in protein-class interactions. An algorithmic strategy could better recover the energetic balance of protein-RNA interactions directly from thermodynamic data. However, the complexity of proteins, combined with the stochastic nature of ROSETTA’s structural refinement workflow, effectively creates a “black box” that provides little information about energy gradients, thereby limiting its ability to optimize structures. These issues, combined with computational expense, make structure optimization inefficient and sometimes infeasible. To this end, we propose using Bayesian Optimization (BO)^12,13^. BO is a derivative-free optimization method that works on expensive-to-compute functions using an uncertainty-aware, easy-to-evaluate surrogate model (e.g., Gaussian Processes, Random Forest, or Bayesian neural networks)^14^. The surrogate model is trained using prior evaluations of the objective function and uses this information to update weight values. These predictions and their respective uncertainties are then fed to an acquisition function that proposes new values to test with the objective function, balancing exploitation (optimizing weights near the optimum) with exploration (sampling unseen regions of the search space). This creates a closed-loop, uncertainty-aware system in which a model continually learns and optimizes.

Here, we developed a closed-loop, experiment-guided strategy for optimizing the weights of the protein-RNA score function within the ROSETTA All-Atom energy framework, using BO as a data-efficient way to explore this parameter space. BO is well-equipped for this, as it handles expensive and poorly defined objective functions, such as estimating energetic contributions at protein-RNA interfaces, by balancing and refining the weights of numerous terms^5^. In the protein field, BO has previously been applied to the regularization of constraints in directed evolution and to the design of functional peptides and antibodies^15–17^. Therefore, we introduce the application of BO to biomolecular complexes, starting with protein-RNA complexes, to establish a design-first technology for predicting optimal protein-RNA interactions and structures.

## Results and Discussion

### Experimental Data Acquisition

We assembled a benchmark of protein-RNA complexes with experimentally measured binding affinities from multiple sources. The training set consisted of 144 non-redundant complexes from Protein-RNA Binding Affinity Benchmark 2.0 (PRBAB) v1.0 and v2.0, spanning diverse protein-RNA interaction classes. The training set includes structurally distinct RNA subclasses, making generalization across a full distribution of interface types a central optimization target for the developed protein-RNA score function. To evaluate generalization, we assembled three independent test sets with no overlap in four-letter PDB identifiers: PDBbind (130 complexes), PRDB v3.0 (66 complexes), and ProNAB (68 complexes)^5,18–20^. We recoded every structure in the dataset into a common two-component (A, B) format to establish consistent energetic decomposition across complexes. Specifically, all protein chains were combined and assigned to chain A, while all RNA chains were merged and assigned to chain B. Partner definitions across datasets were standardized to allow the direct comparison of binding energetics and score functions. Structures acquired from the PDB were set as the “native” conformation.

### Baseline Score Function Performance

We first qualitatively examine the shortcomings of baseline ROSETTA score functions in their ability to capture protein-RNA interactions. Starting from an experimental protein-RNA crystal structure, the Roquin ROQ domain was separated from the RNA and then re-docked using the widely used ref2015 scoring function. The model was then aligned with the original crystal structure (PDB: 5F5H). This revealed that key residues involved in protein-RNA interactions were modeled inaccurately, as shown in **Figure 1**.

**Figure 1.**
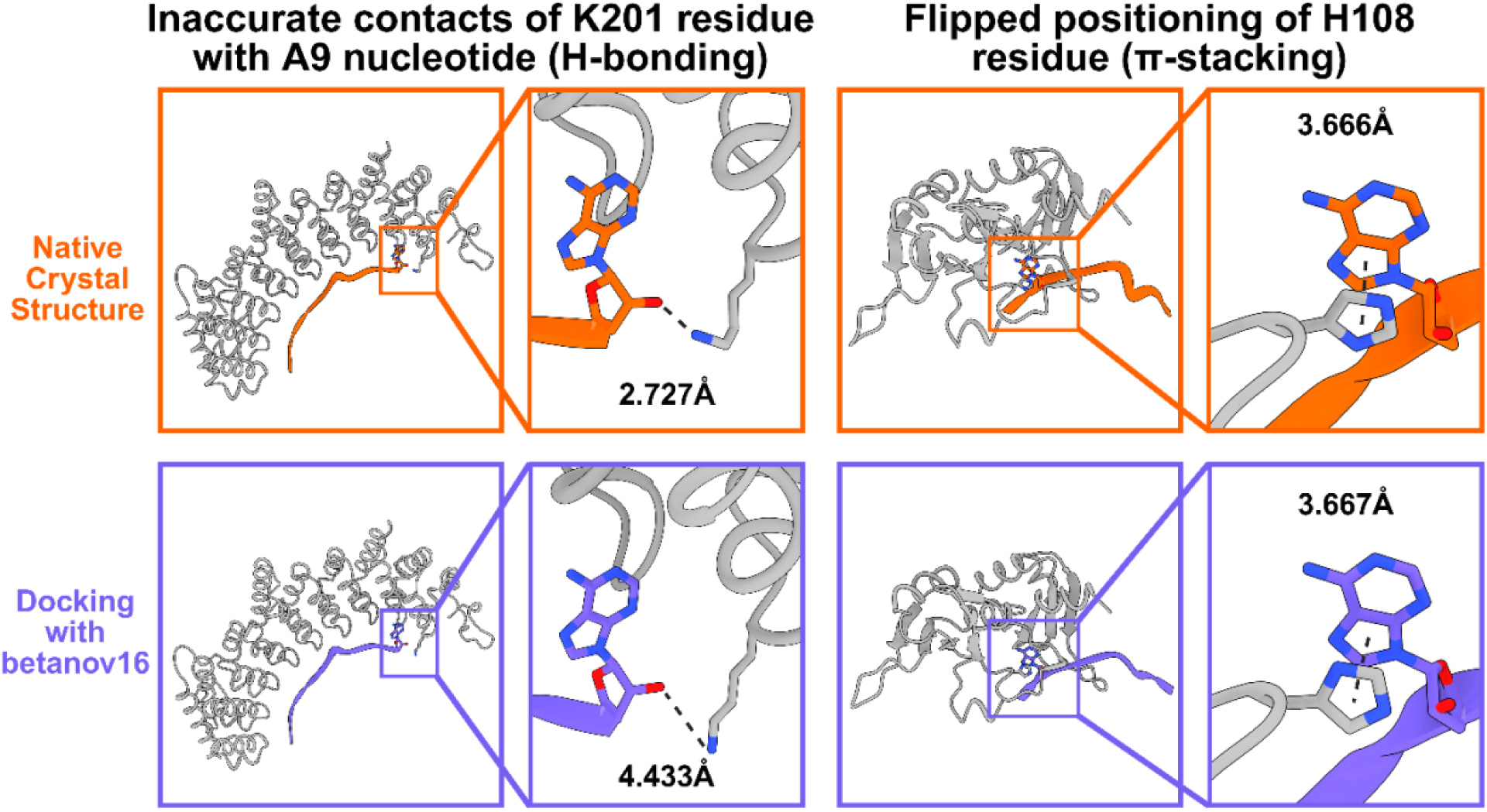
Comparison of protein-RNA crystal structures with a model docked with the betanov16 score function shows inadequate prediction of many key residues involved in protein-RNA interactions, leading to improper hydrogen bond formation in the *Caenorhabditis elegans* PUF protein-RNA complex (PDB: 3QGC)^21^ and π-π stacking in human hnRNP A2/B1 (PDB: 5HO4)^22^.

The ROSETTA all-atom function maintains an approximate 1:1 REU-to-kcal/mol conversion under the condition that ratios between each score function term are held constant^10^. However, protein-RNA interfaces are more driven by electrostatics than protein-protein interfaces, and as a result, protein-trained scoring functions are not designed to capture the physicochemical driving forces of the protein-RNA binding surface^5,23^. Based on the claimed conversion, we reasoned that the difference between the predicted REU and the experimental kcal/mol would be a reasonable metric for comparing score functions.

To quantify the observed errors, we benchmarked baseline ROSETTA score functions spanning protein- and RNA-centric parameterization on a standardized set of protein-RNA complexes with known experimental binding free energies. We evaluated the best-performing protein- and RNA-trained all-atom functions (betanov16 and RNA hires, respectively) across the training set and test sets. Baseline score functions produced wide error distributions with noticeable variability in both overall spread and median bias across datasets (**Figure 2**).

**Figure 2.**
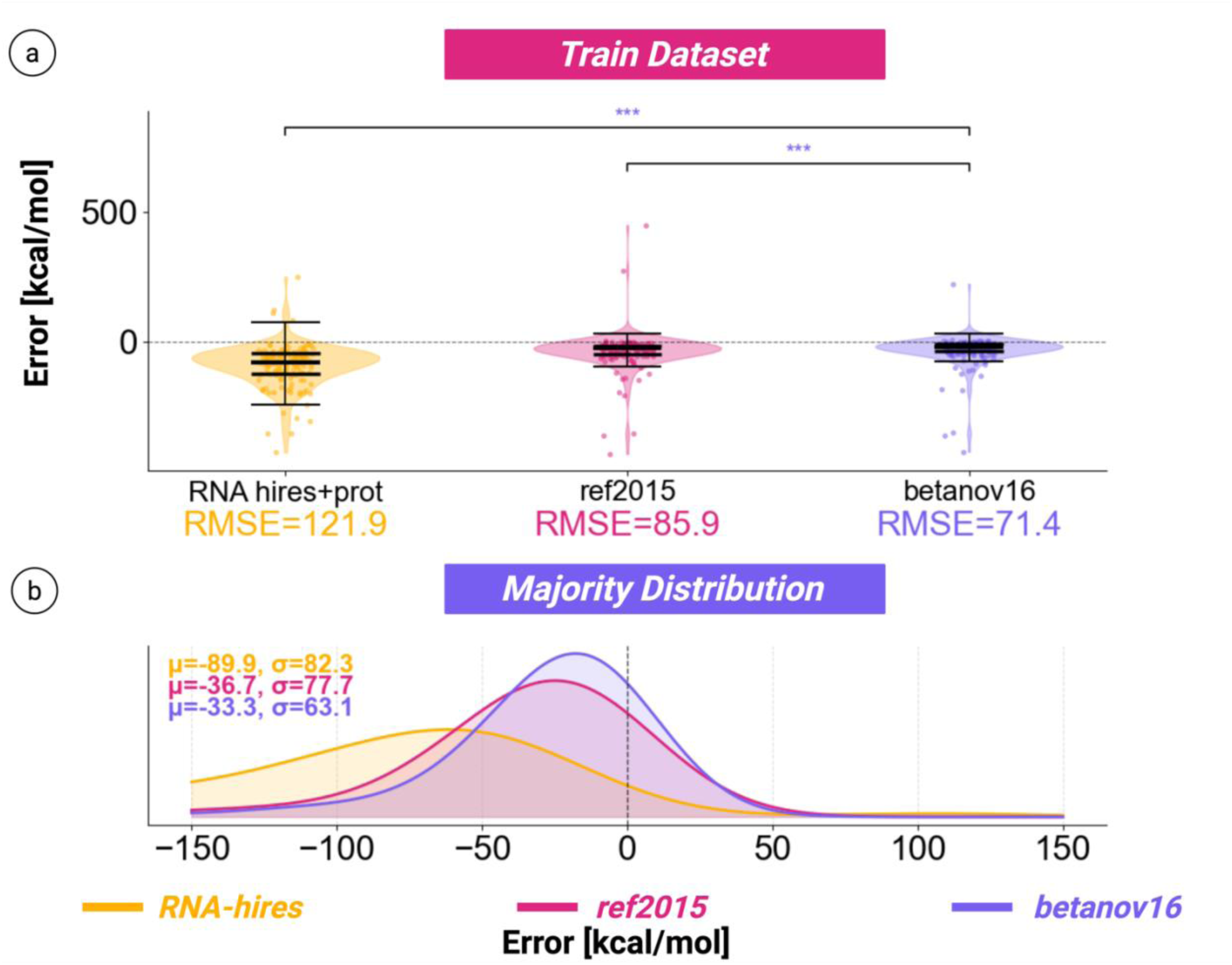
Error across baseline ROSETTA score functions. (A) highlights the RMSE of each of the out-of-the-box score functions, while (B) displays the average error of each, alongside the standard deviations. Distributions were made using a kernel density estimator. We find that betanov16’s improvement over the other two score functions was statistically significant, as determined by a Wilcoxon Signed-Rank test.

The betanov16 score function exhibited a consistent negative bias relative to experimental binding free energies, whereas RNA hires often produced even larger deviations and greater dispersion. Prediction accuracy also varied with complex size and interface composition, implying that neither protein-centric nor RNA-centric parameterizations fully capture binding energetics governing protein-RNA binding within a simple bound-bound model.

### Finetuning betanov16 using Bayesian Optimization

We sought to refine the top protein-protein all-atom energy function by adjusting only its most important terms. We selected *betanov16* as the baseline score function for optimization because it exhibited the strongest agreement with experimental protein-RNA binding affinities among the ROSETTA score functions tested. Protein-RNA recognition is determined by a distinct interface between the two partners rather than by the global folding behavior of either chain alone; specifically, electrostatic forces, rather than hydrophobic contributions, drive protein-protein binding. To maintain the global accuracy of each individual chain in the protein-RNA complex and to address the established limitation that existing score functions lack the physical grounding to generalize across the structural diversity of RNA classes without targeted reweighting of intermolecular terms, we only allowed 9 (out of 26) score function terms to change during optimization. The nine terms chosen for optimization are dominant contributors to the intermolecular binding energetics and geometry at protein-RNA interfaces, as revealed by structural analysis of the PRBAB database^5^. These forces include long- and short-range backbone interactions, hydrogen bonding, solvation, van der Waals contacts, and electrostatic interactions. We kept the remaining 17 *betanov16* global terms, such as rotamer preferences, backbone torsion penalties, and disulfide geometry, constant. Each term was assigned an adjustable bound during optimization.

Our strategy followed the approach demonstrated by Shringari *et al*., who showed that non-linear reweighting of ROSETTA energy terms against experimental values yields a custom score function with improved predictive accuracy over an unmodified baseline without requiring additional score-function terms^11^. Conservative search bounds were set from the range of values present in baseline ROSETTA score function variants. This methodology follows the strategy in the ROSETTA all-atom framework where individual terms are adjusted for specific interactions, such as the upweighting of disulfide geometry, and the modulation of hydrogen bond and solvation terms across different molecular systems, showing that application-specific reweighting is physically valid within the ROSETTA framework^10^. Optimization bounds were set to preserve as much of the 1:1 REU-to-kcal/mol relationship as possible while still allowing rebalancing of interface contribution terms.

By constraining optimization to known influential parameters, we prevent wasted evaluations on unimpactful weights, improving computational efficiency. In addition, we prevent the natural balance inherent to *betanov16.* Search bounds for each term were set using ranges in the current ROSETTA score functions, ensuring physically sensible limits while preserving sufficient flexibility for energetic reweighting^5,6,24^. We set the computed total error (*E_t_*_otal_) as our objective function for BO, as shown in **Figure 3**, where *ΔG_i_* and *ΔĜ_ι_* are the true and predicted energies, respectively, as given in Equation (1). The RMSE is computed on the predicted dataset to quantify total error, acknowledging that reweighting individual intermolecular terms “breaks” the fixed-ratio condition; however, this trade-off was necessary to maintain consistency in the comparison metric.

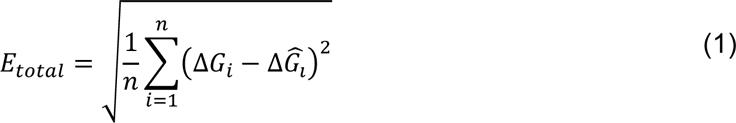

**Figure 3.**
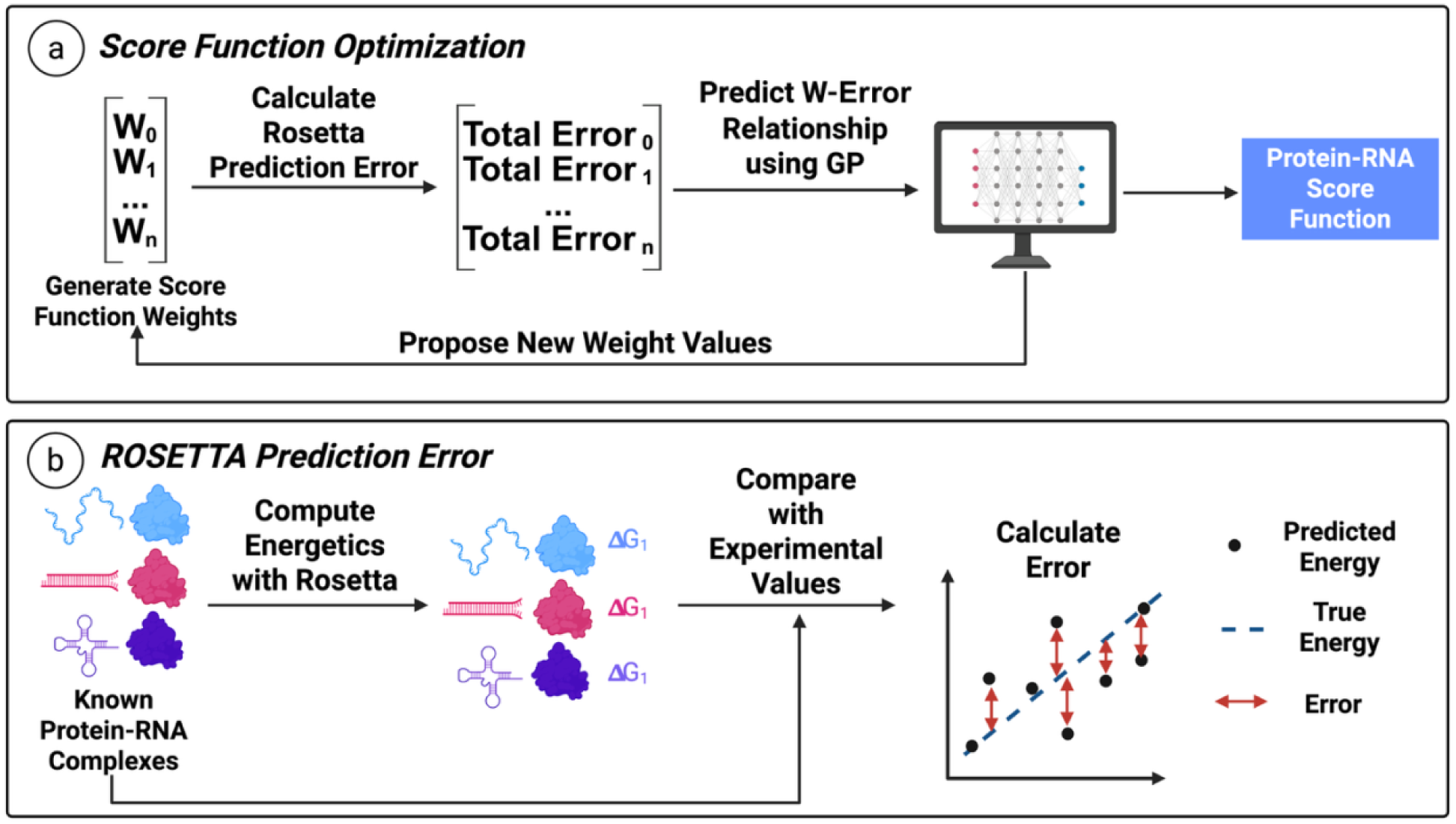
Bayesian Optimization for Score Function Tuning Overview. (A) A subset of protein-RNA complexes with experimentally validated energies was used to (B) train Rosetta to predict energy values. Total error was determined from these predictions. The total error was minimized using BO, in which the relationship between the provided score function weights and errors was learned via a GP model. These predictions informed subsequent weights, which were tested on new, unseen data.

We used a Gaussian process (GP) model to learn the relationship between provided ROSETTA weights and *E_total_*, due to its fast-to-compute and uncertainty-aware nature. The GP-predicted values and uncertainties were provided to an acquisition function to propose new weights to test in the ROSETTA protocol (**Figure 3**). To implement BO, we used *BoTorch,* including a Matérn 5/2 covariance kernel for the GP and Expected Improvement for the acquisition function^25,26^. To initialize our campaigns, we used scrambled Sobol sequences to improve the diversity of starting information. Each campaign was provided the same seed, ensuring uniform starts.

### Bayesian Optimization Results

Across five independent optimization runs (10 initial Sobol samples, 30 iterations, 40 total), the objective dropped quickly in the early steps and leveled off to a stable plateau around 15 trials (**Figure 4**). As a baseline, we included a random search method. This method removes the acquisition process, representing a completely blind, surrogate-agnostic model. Five separate campaigns were run, each with a unique set of initializations. These initial weights were made consistent between the random search and BO. The random search, evaluated under the same search space and objective function, showed slower improvement, greater inter-run variability, and a higher final objective value (**Figure 4**; grey). This pattern indicated that BO efficiently identifies productive score-term reweighting within a limited evaluation budget and that convergence cannot be attributed to random parameter exploration.

**Figure 4.**
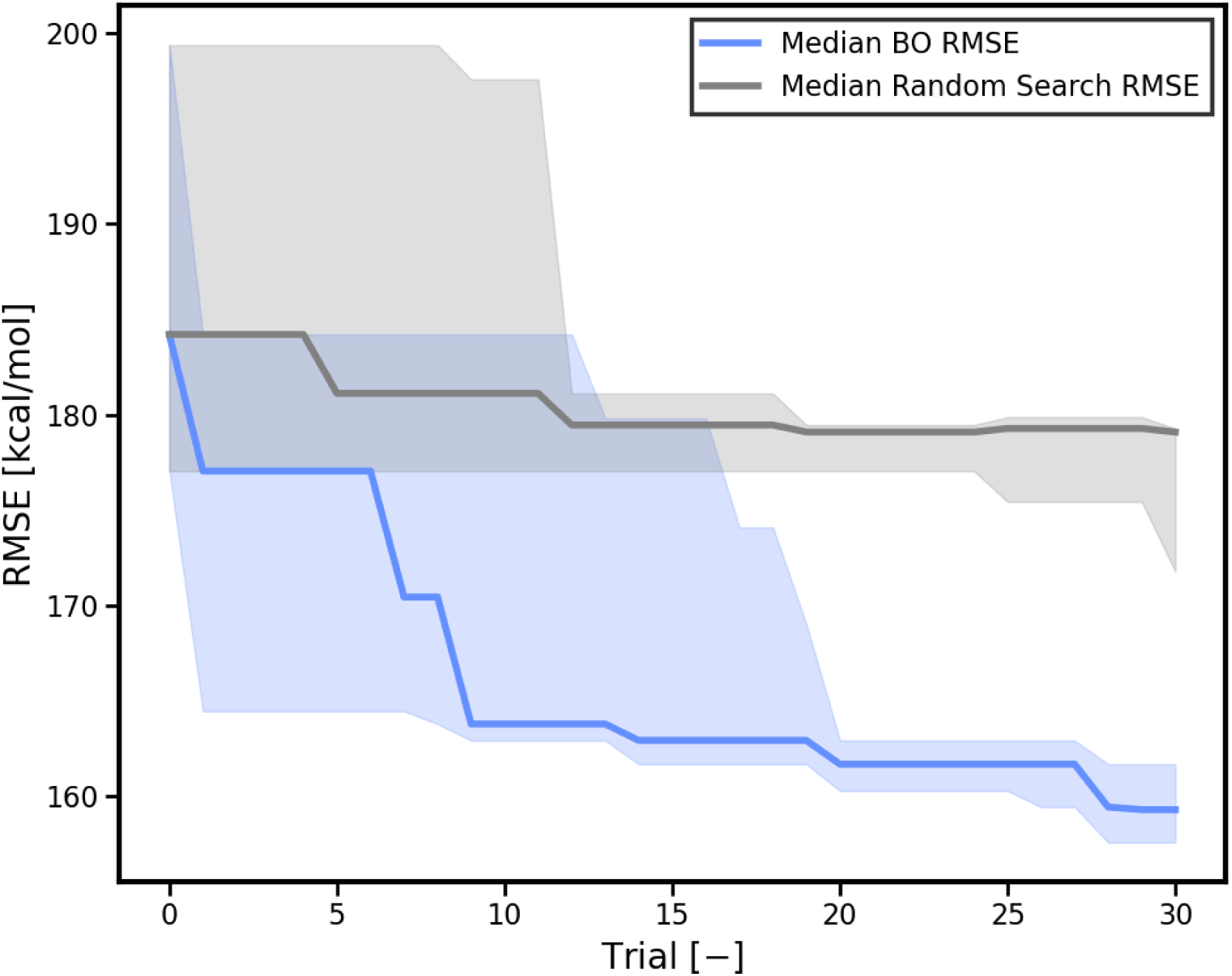
Lowest RMSE by trial of BO campaigns against Random Search. BO demonstrates the ability to find a better set of score function weight values in fewer computations than random search. We minimized root mean squared error (RMSE) between the predicted ΔG_bind_ (REU) and experimental ΔG_exp_ (kcal/mol). The qualitative alignment between known and predicted structures demonstrates this improved convergence.

### Analysis of New Weight Values

Results of the Bayesian Optimization protein-RNA (boRNA26) re-weighting are provided in **Table 1**. The nine terms chosen for optimization include van der Waals (attractive and repulsive contributions), solvation, electrostatic interactions, hydrogen bonding (short- and long-range backbone-backbone interactions, backbone-sidechain, and sidechain-sidechain interactions), and other orientation-dependent polar interactions. Backbone-sidechain, sidechain-sidechain, implicit solvation, and orientation-dependent solvation weights increased, while attractive and repulsive forces, short- and long-range backbone hydrogen-bonding, and electrostatics decreased. These shifts seem to suggest that baseline protein-trained score functions tend to “overweigh” polar and charged interactions when applied to protein-RNA interfaces. This over- stabilization likely stems from compensating for heavily charged backbones and a dense network of potential hydrogen-bonding sites. These properties can cause static models to overstate the strength of polar interactions, in turn driving predicted binding energies into less favorable ranges.

**Table 1.** Comparison of Energy Terms: betanov16 vs. boRNA26.

| Interaction class | Score Term | Description | betanov16 | boRNA26 | % Change |
| --- | --- | --- | --- | --- | --- |
| *van der Wals/Packing* | fa_atr | Attractive energy (*dist. d*) | 1.00 | 0.7 | -30% |
|  | fa_rep | Repulsive energy (*dist. d*) | 0.55 | 0.40 | -27% |
| *Solvation* | fa_sol | Gaussian exclusion implicit | 1.00 | 1.24 | +24% |
|  | lk_ball | Orientation-dependent polar | 0.92 | 1.30 | +41% |
| *Electrostatics* | fa_elec | Interaction between charges | 1.00 | 0.50 | -50% |
| *Hydrogen bonding* | hbond_sr_bb | Short-range backbone | 1.00 | 0.6 | -60% |
|  | hbond_lr_bb | Long-range backbone | 1.00 | 0.6 | -60% |
|  | hbond_bb_sc | Backbone-side-chain | 1.00 | 1.6 | +60% |
|  | hbond_sc | Side-chain-side-chain | 1.00 | 1.6 | +60% |

Several observations support this interpretation. First, short- and long-range backbone hydrogen-bond weights decreased, while backbone sidechain hydrogen-bond weights increased, emphasizing a global energetic imbalance rather than isolated errors within protein hydrogen - bonding subcategories. Second, both the implicit and orientation-dependent solvation terms increased, which fits the idea that ordered water molecules at protein-RNA interfaces carry a real energetic cost. The increase in the solvation score-term weight helps capture the cost of disrupting hydration networks and the stabilizing role they play within the binding pocket. Lastly, global scoring offsets move in a consistent direction as the optimization converges, suggesting that the main problem with the baseline scoring functions is not missing interactions but the relative scaling of those already present, particularly for protein-RNA interactions.

### Score Function Comparison

Although boRNA26 REU values are no longer directly comparable to kcal/mol due to this reweighting, optimization yields closer agreement with experimental ΔG values than either betanov16 or ref2015 across all test sets, and structural predictions qualitatively improve key interaction geometries (**Figure 5**). The energetic signal between predicted and experimental values is preserved and physically grounded even where the strict unit mapping is not. The improvement in both prediction accuracy and structural penalty substantiates our reweighting approach.

**Figure 5.**
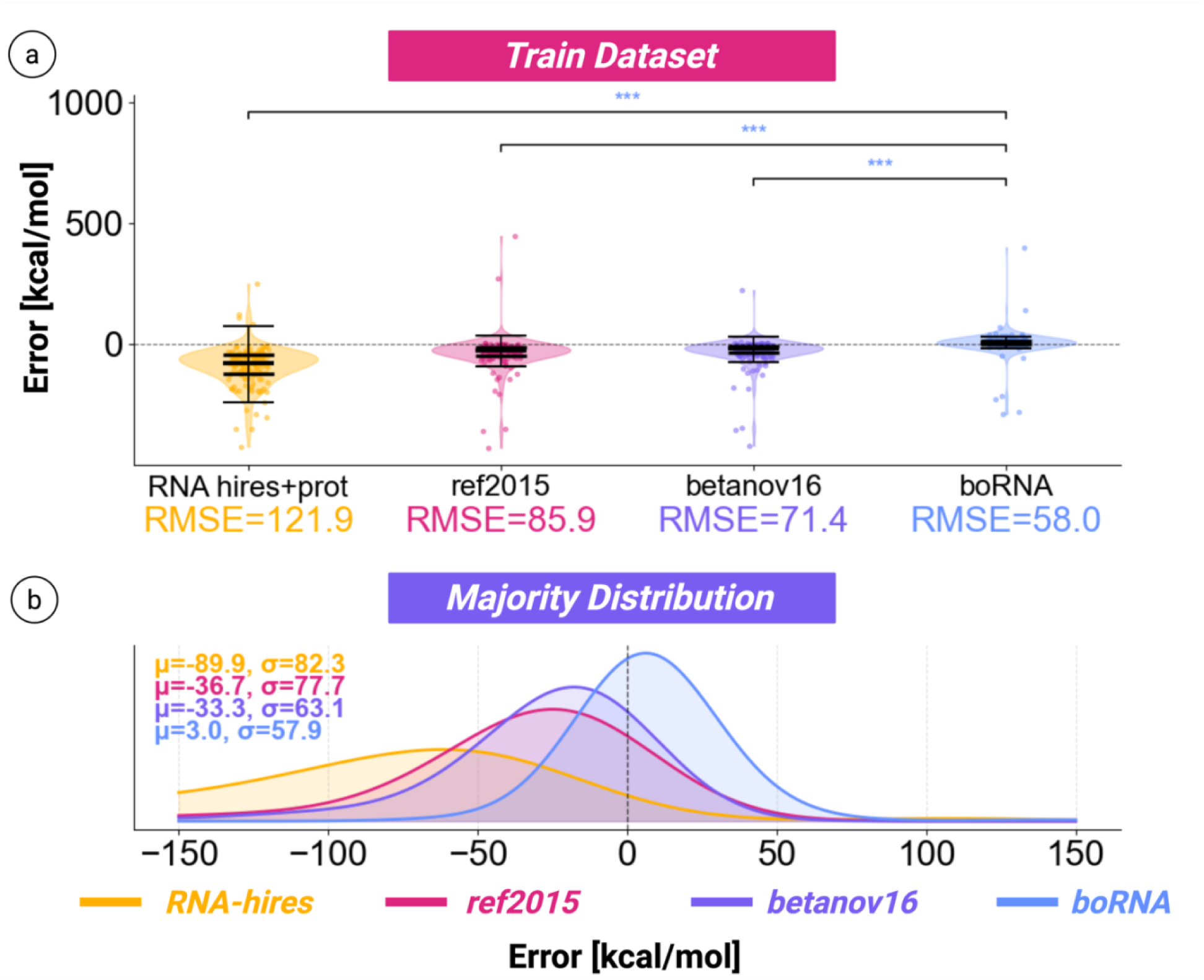
Comparisons between ref2015, betanov16 and boRNA (BO-tuned score function). (A) RMSE error used to evaluate differences between the score functions, where statistical significance was determined using the Wilcoxon-Signed-Ranked test. The improvement of boRNA over baseline score functions was statistically significant. (B) Outliers were removed, and the distribution was smoothed using a kernel density estimate to showcase boRNA’s improved mean and consistency (*σ*).

### Score Function Generalization Across Independent Datasets

To evaluate generalization, we assembled three independent test sets with no overlap in four-letter PDB identifiers: PDBbind (130 complexes), PRDB v3.0 (66 complexes), and ProNAB (68 complexes)^5,18–20^ (**Figure 6**), where we evaluated comparisons of docking across π-π stacking, ribose bonding, nucleobase contacts, and phosphate backbone bonding. These docking studies demonstrate that boRNA26 can achieve tighter alignment with native crystal structures than existing scoring functions. Using these data, we sought to improve the scoring function’s predictive capacity. Therefore, we first ensured that the data found in these sets were independent of one another to prevent any leakage (**Figure 7A**). **Figure 7B** shows that the performance of boRNA is not limited to just one model. Although the variance increases when using boRNA, the mean absolute error remains the smallest across all test sets, highlighting the improved performance of reweighting. We highlight multi-objective optimization, in which both error and variance are minimized, as a potential avenue for future expansion^8,13^.

**Figure 6.**
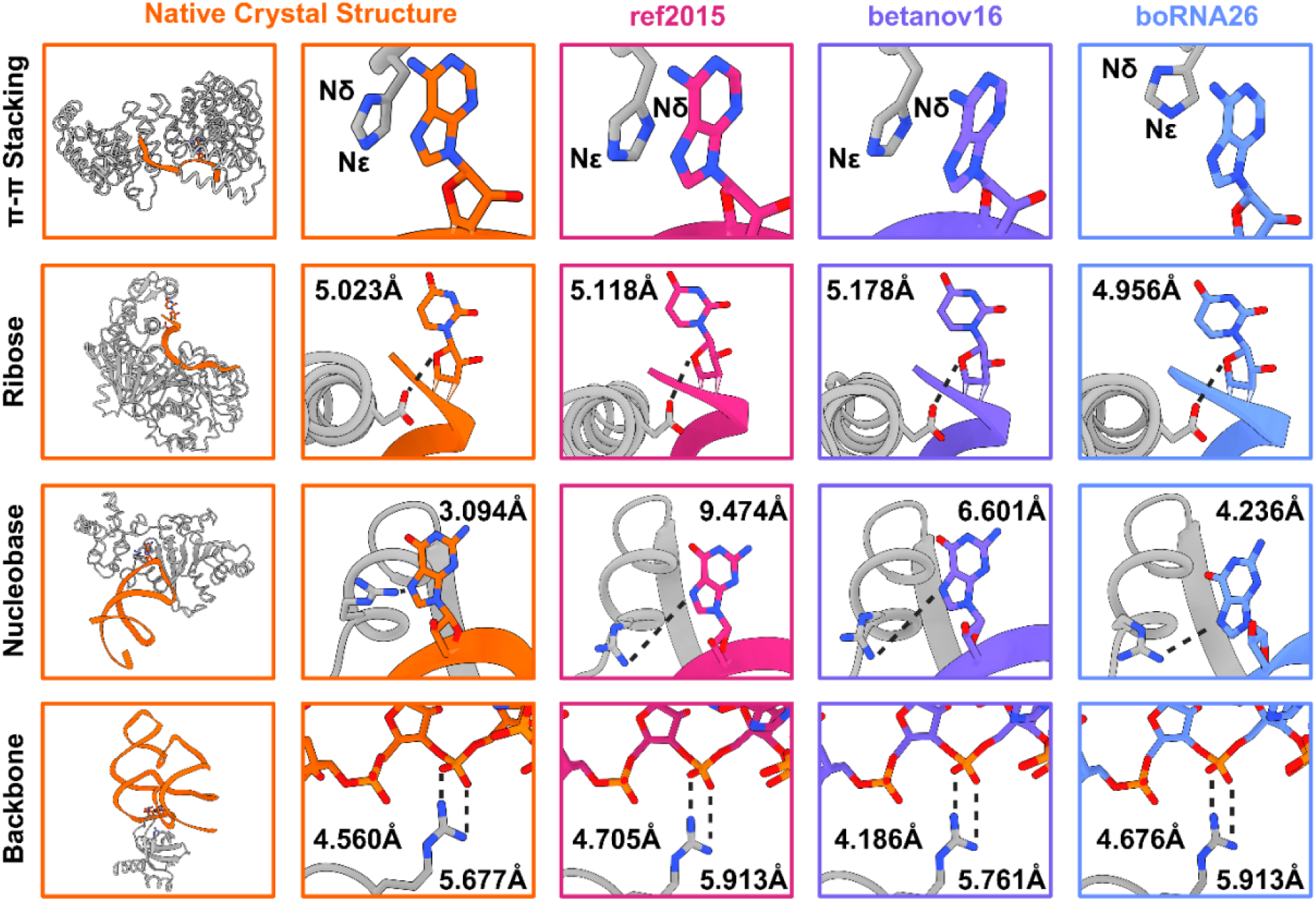
Docking with a BO-tuned score function. Comparisons of docking with the betanov16 and ref2015 score functions can result in π-π stacking (PDB: 5WTY)^27^, ribose bonding (PDB: 3I5X)^28^, nucleobase contacts (PDB: 6AAX)^29^, and phosphate backbone bonding (PDB: 7KKV)^30^ that better align with the experimental structure of a protein-RNA complex.

**Figure 7.**
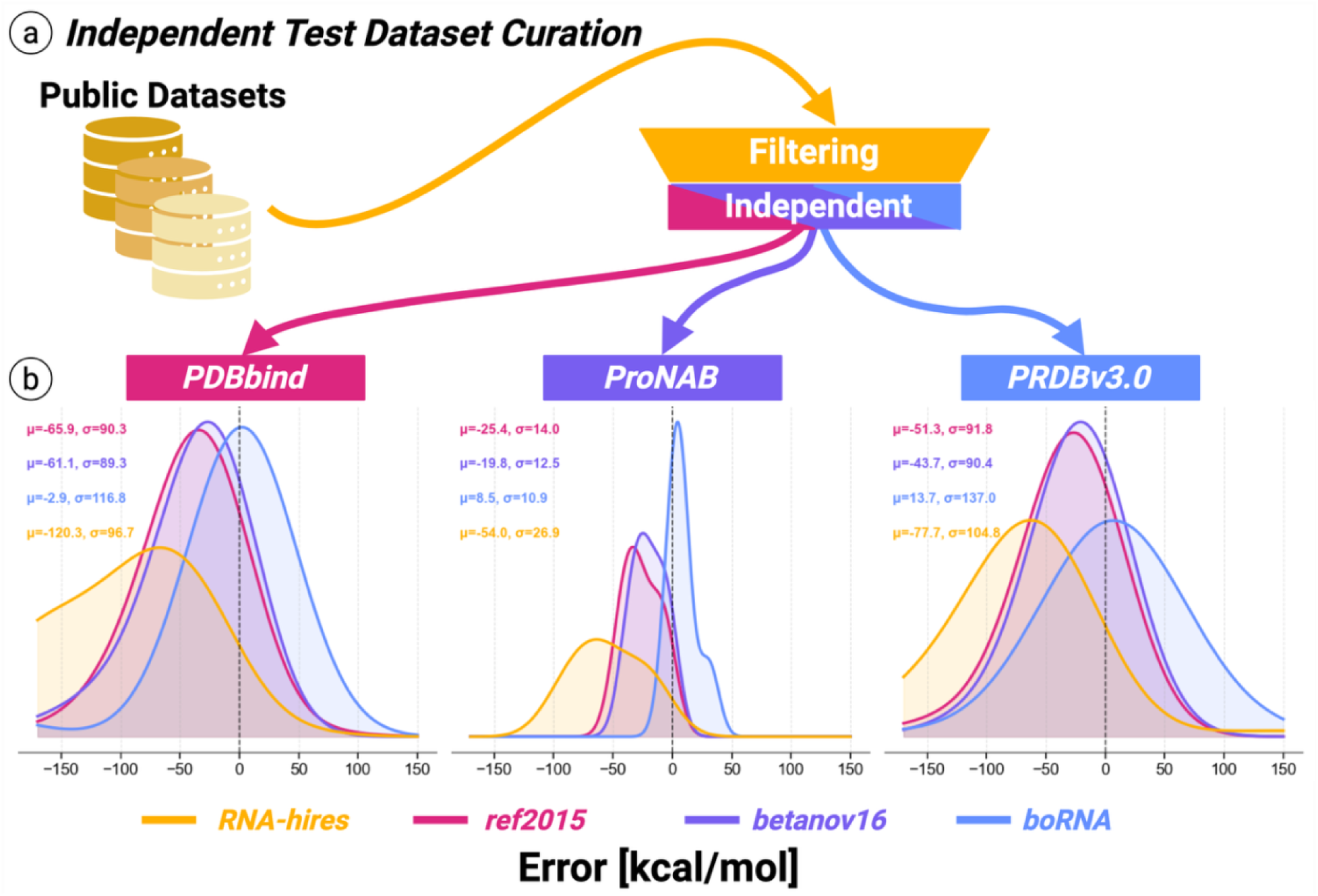
Improving predictive capacity via database filtering and optimization. (A) Online datasets were acquired and filtered to remove redundant and nonconverging structures. This process resulted in 3 test sets, all independent of one another. (B) boRNA shows a lower average error relative to betanov16 and ref2015 across all 3 datasets, highlighting its improved predictive capacity.

### RNA-Protein Subtype Analysis

RNA is a diverse class of biomolecules with subtypes that differ in structure, bonding networks, and interactions. Based on interface geometry “signatures”, RNA binding proteins (RBPs) have specific interactions with these RNA subclasses (**Table 2**). These signatures result in quantitative and qualitative differences, and thus variations in total score functions, between functional classes. Because of these subtle differences, we hypothesized that baseline ROSETTA protein-protein trained score functions will misallocate weights across RNA subclasses compared with the protein-RNA optimized weight set (boRNA). Additionally, a score function tuned to a specific class would not capture global protein-RNA trends or the geometric subtleties of other subclasses. Results show that boRNA outperformed ref2015 and betanov16 across all tested RNA subtypes, with the largest specific improvements for rRNA, tRNA, and snRNA, and a global improvement for both single- and double-stranded (ssRNA and dsRNA) protein-RNA complexes (**Table 3**).

**Table 2.**
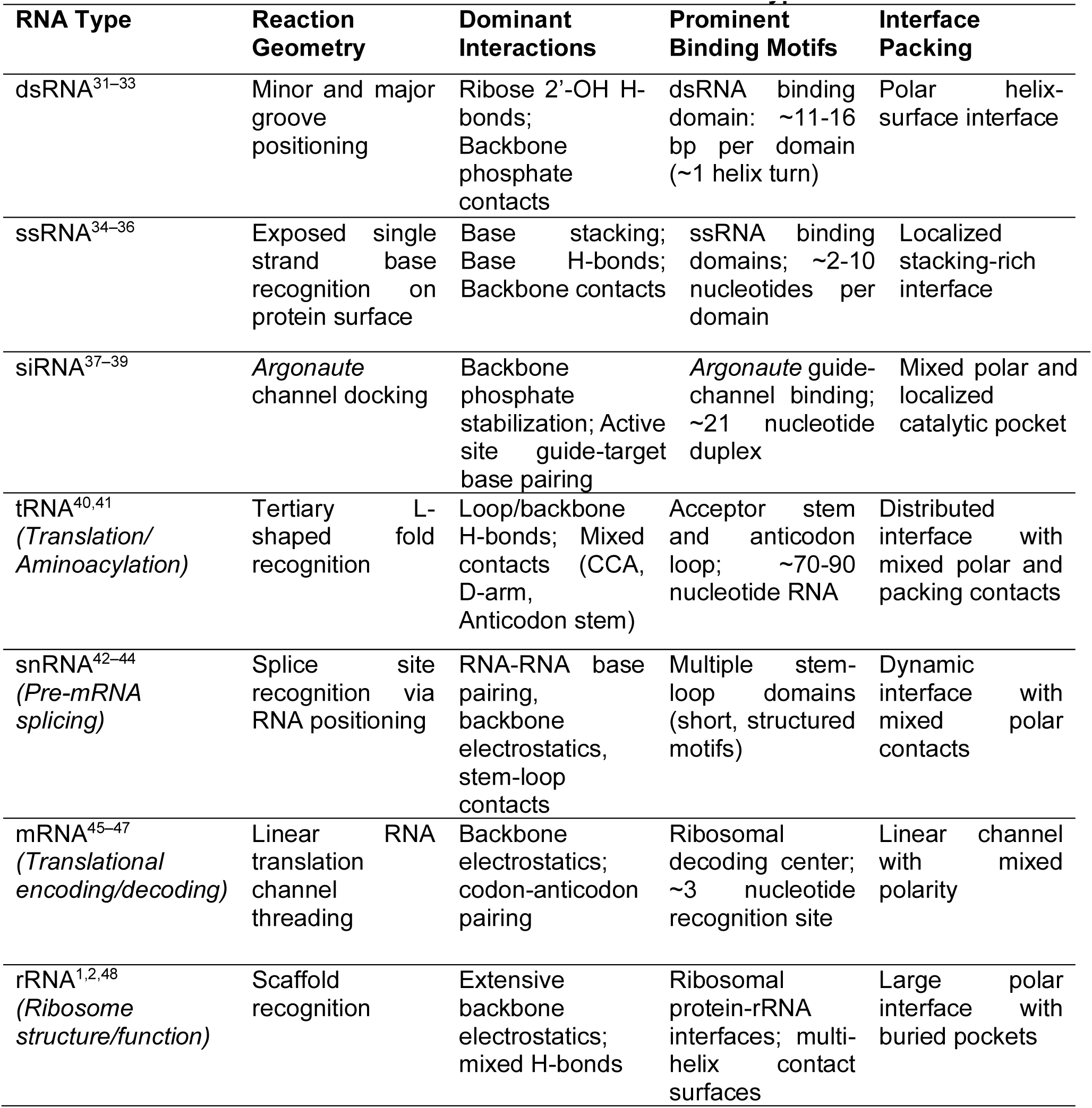
Differences in Protein-RNA Interactions Across RNA Subtypes.

**Table 3.** Error ± Standard deviation of tested score functions on RNA subclasses.

| RNA Type | boRNA<br>Error ± Std | ref2015<br>Error ± Std | betanov16<br>Error ± Std |
| --- | --- | --- | --- |
| dsRNA | 10.85 ± 8.62 | -18.72 ± 16.02 | -9.75 ± 13.86 |
| ssRNA | 3.47 ± 10.59 | -32.30 ± 23.40 | -26.26 ± 20.00 |
| siRNA | 9.17 ± 2.01 | -12.99 ± 7.26 | -15.25 ± 17.64 |
| tRNA | 2.07 ± 7.17 | -43.52 ± 23.35 | -33.15 ± 19.11 |
| snRNA | -1.90 ± 15.23 | -85.55 ± 70.72 | -78.35 ± 66.24 |
| mRNA | 7.47 ± 12.65 | -26.77 ± 11.22 | -22.34 ± 14.97 |
| rRNA | 4.73 ± 5.80 | -33.14 ± 18.32 | -24.97 ± 19.07 |

Sore function accuracy improvements correspond with the weight shifts described above. Outlier structures that failed to converge were removed using the interquartile range (IQR) method prior to comparison. Among the subclasses we tested, tRNA, which has distributed interface contacts across loops, backbone, and a mixed polar and packing character, was scored most accurately with increased backbone-sidechain and sidechain-sidechain hydrogen-bonding weights. The increase in the solvation term also benefited rRNA scoring, consistent with its buried polar interfacial pockets. snRNA showed the largest overall improvement, which may reflect the reduced van der Waals and packing weights better accounting for its short, tightly structured stem-loop motifs. Electrostatic-dominated interfaces, such as dsRNA, are captured less completely by these adjustments, consistent with decreased backbone hydrogen-bonding and reduced weightings of the electrostatic term, resulting in a modest gain in accuracy.

## Conclusion

Here, we show that Bayesian optimization can be used to improve the agreement between ROSETTA predictions and experimental data when scoring protein-RNA interactions. We found that our BO score function outperformed betanov16 when trained and tested on independent data sets. Here, we proposed a fine-tuned ROSETTA score function that robustly handles protein-RNA interactions regardless of experimental origin. Through this, we generated a score function that accurately captures significant RNA subclass-specific interactions, thereby validating its physical accuracy. Further, we establish Bayesian Optimization as a novel approach to directly optimize ROSETTA score functions for any protein-class interaction. Our score function establishes high confidence that *in silico* protein-RNA design increases, as physical and experimental agreement improves. Unlike prior ROSETTA score-function studies, which train machine-learning models to predict values, our BO method directly improves the underlying ROSETTA BEHAVIOR. This study shows that BO can be used for fine-tuning biomolecular score functions to better reflect experimental data. Future studies are expected to extend this program to account for the important roles that ligands and small molecules^49^, engineered protein binders ^8,50^, amino acid and nucleobase mutations^51^, and metal co-factors^52,53^ may play in governing protein-RNA interactions, enabling a tunable space for directly designing and optimizing novel protein-RNA complexes.

## Methods

### Score Function Optimization

Weight optimization was done using Bayesian optimization (BO) applied to nine energy terms in the ROSETTA betanov16 score function. The weighted combination of these terms determines specific interactions at protein–RNA interfaces: Lennard-Jones attraction and repulsion (fa_atr, fa_rep), implicit solvation and orientation-dependent hydration (fa_sol, lk_ball), electrostatics (fa_elec), and four hydrogen bonding terms spanning backbone-backbone, backbone-sidechain, and sidechain-sidechain contacts (hbond_sr_bb, hbond_lr_bb, hbond_bb_sc, hbond_sc. All other terms were held fixed at their betanov16 default values. Search bounds for each optimized term were set to explore around betanov16 defaults, allowing variations of approximately ±30-60% depending on the term. For example, fa_elec was allowed to range from 0.50 to 1.60 (betanov16 default: 1.0), while fa_atr was bounded between 0.70 and 1.35 (betanov16 default: 1.0). Bounds were chosen to span the range of published ROSETTA protein and RNA score function variants while excluding physically implausible weight combinations.

### Structural Dataset and Preparation

Optimization was done using 143 protein-RNA structures from the protein–RNA binding affinity benchmark (PRBAB) v2.0. Each complex was paired with an experimentally measured binding free energy (ΔG_exp_, kcal/mol). For NMR ensembles, only the first structure was used. All structures were initially cleaned of ligands, metal ions, and water molecules and assigned a uniform two-chain representation: chain A represents the protein, and chain B represents the RNA.

### Binding Energy Calculation

For each candidate weight set, the predicted binding energy was calculated in Equation (2) as:

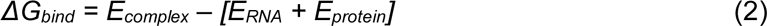

where each component was scored independently using the same trial score function, isolating interfacial contributions from intramolecular terms.

#### Optimization objective

We minimized root mean squared error (RMSE) between the predicted ΔG_bind_ (REU) and experimental ΔG_exp_ (kcal/mol) across all 143 structures. No explicit unit conversion was applied between REU and kcal/mol. The optimizer was free to implicitly rescale via weight-magnitude adjustments.

### Bayesian Optimization Campaign

BO campaigns were initiated with scrambled Sobol samples. Each campaign started with 10 of these data points, determined by the campaign number. This method allowed us to demonstrate the ability for BO to converge on better weight values across distinct sets of starting points. The campaign was run for 30 iterations of the surrogate model-acquisition function selection process, resulting in 40 points evaluated altogether for each campaign. The random search removes the acquisition function and instead randomly selects an unexplored point.

### Gaussian Processes

For BO, we used GPs as our surrogate model, although other methods, such as Bayesian neural networks and random forests, can also be used. We chose GPs because they are relatively simple to compute compared to the ROSETTA function and, by nature, have accessible uncertainty quantification for the acquisition function. GPs are nonparametric, setting a distribution across potential functions *Pr*(*f*(*x*)|*θ*), where *θ* represents the data points observed thus far. This distribution is characterized by mean *μ* and covariance Σ, which is itself obtained via a kernel function *κ*. We set a *μ* to 0, scaling our training data accordingly. For boRNA, we used the Matern 5/2 kernel, which is a standard in the field, defined in Equation (3) as:

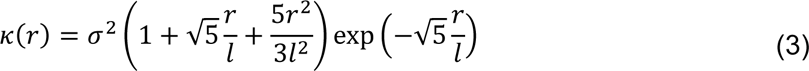

Where *r* is the 2-norm between two datapoints and *l* is the length scale hyperparameter, which is fitted for each of the score function weights modified.

### Acquisition Function

We used expected improvement (EI) as the acquisition function of choice, defined in Equation (4):

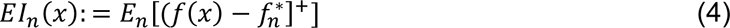

In non-batch systems, we select the point that maximizes the probability of improvement upon observation. Since we are using a GP, assuming the functions are normally distributed yields a Gaussian posterior, and thus EI yields a tractable, closed-form expression that we maximize directly. Alternatives include upper confidence bound (UCB) and Thompson sampling.

## Acknowledgements

This work was supported in part by The Ohio State University Center for Cancer Engineering, Curing Cancer Through Research in Engineering and Sciences, and the National Cancer Institute (NCI) through the R21 grant R21CA312456. B.R.K. acknowledges financial support from the Prostate Cancer Foundation Young Investigator Award. This work used the Ohio Supercomputer Center, which provides High Performance Computing resources and expertise to academic researchers across the State of Ohio. OSC is a member of the Ohio Technology Consortium, a division of the Ohio Department of Higher Education. Figures have been created with UCSF ChimeraX and Biorender.com.

## Author Contributions

Joseph Bailey: Led idea generation for the work. Wrote the original draft of the manuscript, generated figures and graphics for the manuscript, edited, and revised the manuscript. Nathan Phan: Supported idea generation for the work, wrote the original draft of the manuscript, generated figures and graphics for the manuscript, edited, and revised the manuscript. Søren Spina: Assisted with writing the original draft of the manuscript, generating figures and graphics for the manuscript, editing, and revising the manuscript. Rachel Getman: Edited and revised the manuscript. Joel Paulson: Supported idea generation for the work, edited and revised the manuscript. Blaise Kimmel: Led idea generation for the work. Wrote the original draft of the manuscript, edited and revised the manuscript, approved the final version of the manuscript, and acquired funding to support the work.

## Data Availability

Datasets, source data, and code are provided on GitHub (Kimmel-Lab).

