## Supplementary material for "Experimentally Tuned Protein-RNA Rosetta Score Function using Bayesian Optimization": SI

**Supplementary Information**

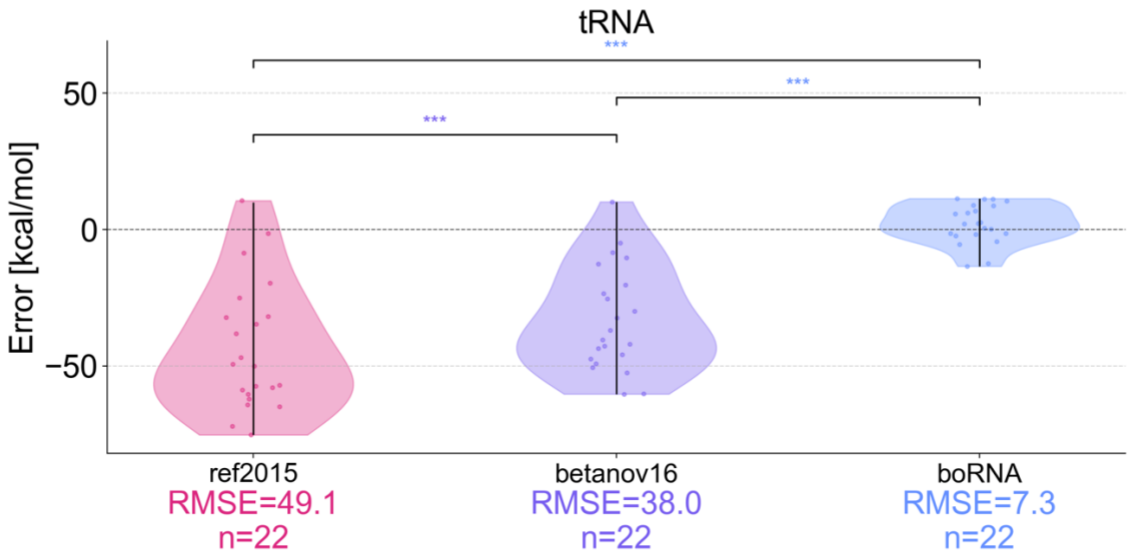


**Supplementary Figure 1.** tRNA Prediction Error.

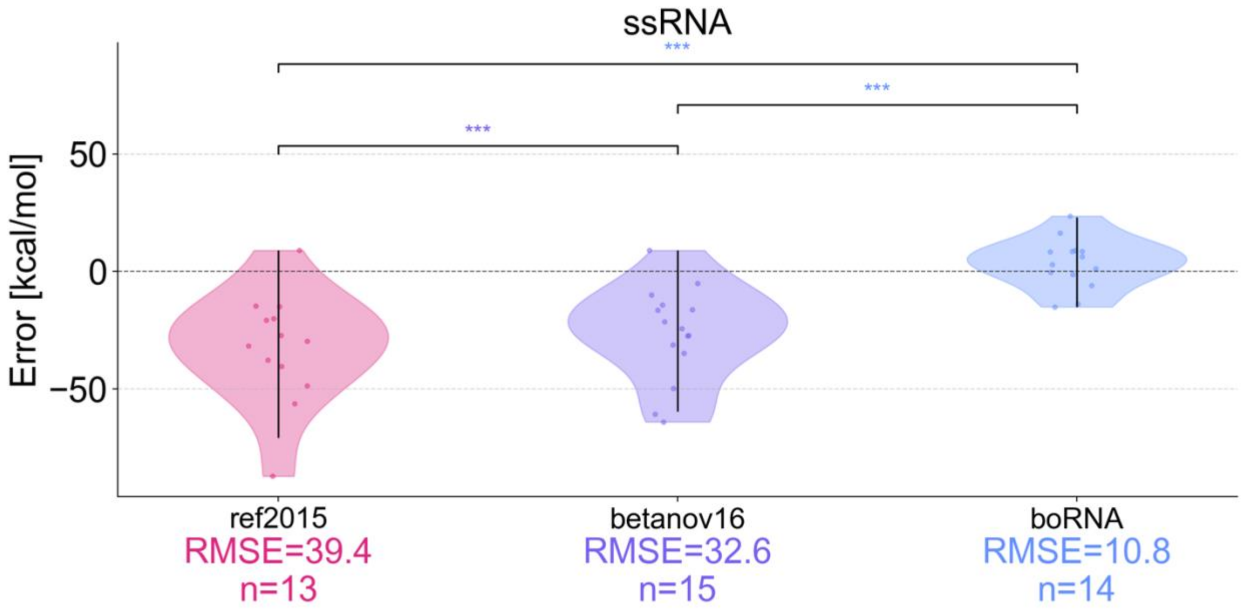


**Supplementary Figure 2.** ssRNA Prediction Error.

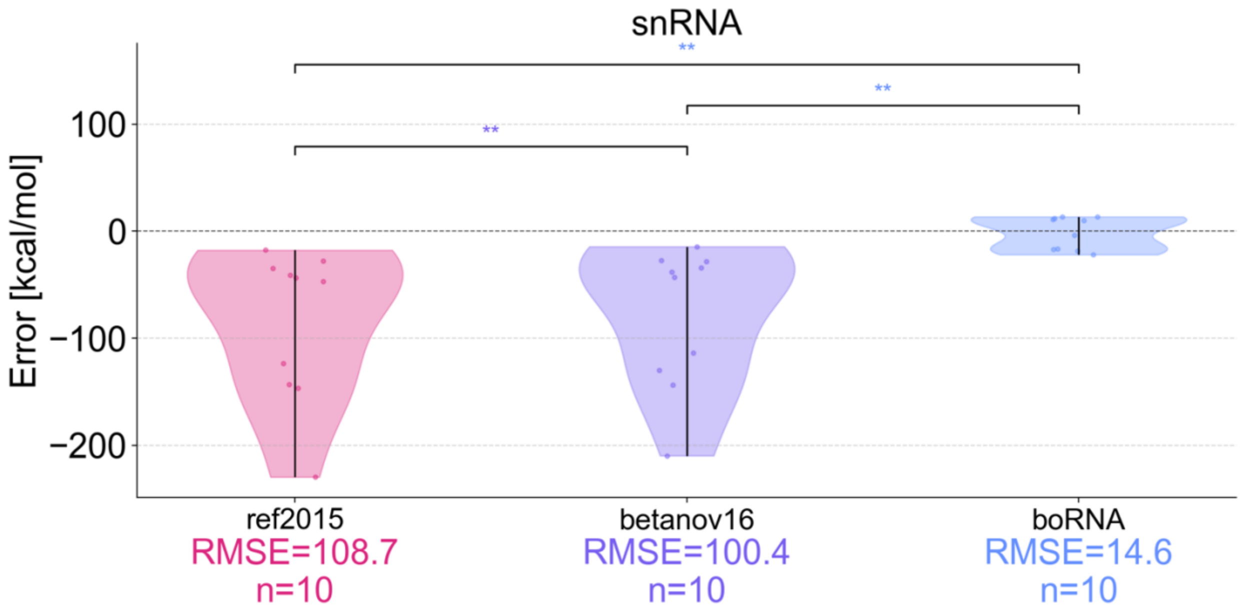


**Supplementary Figure 3.** snRNA Prediction Error.

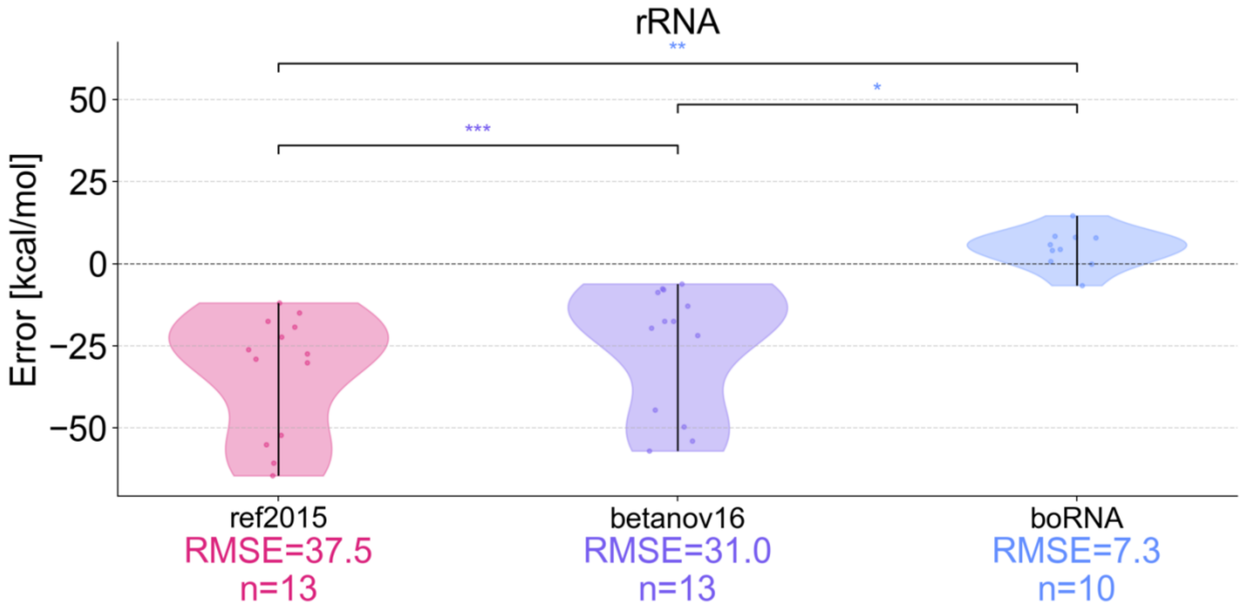


**Supplementary Figure 4.** rRNA Prediction Error.

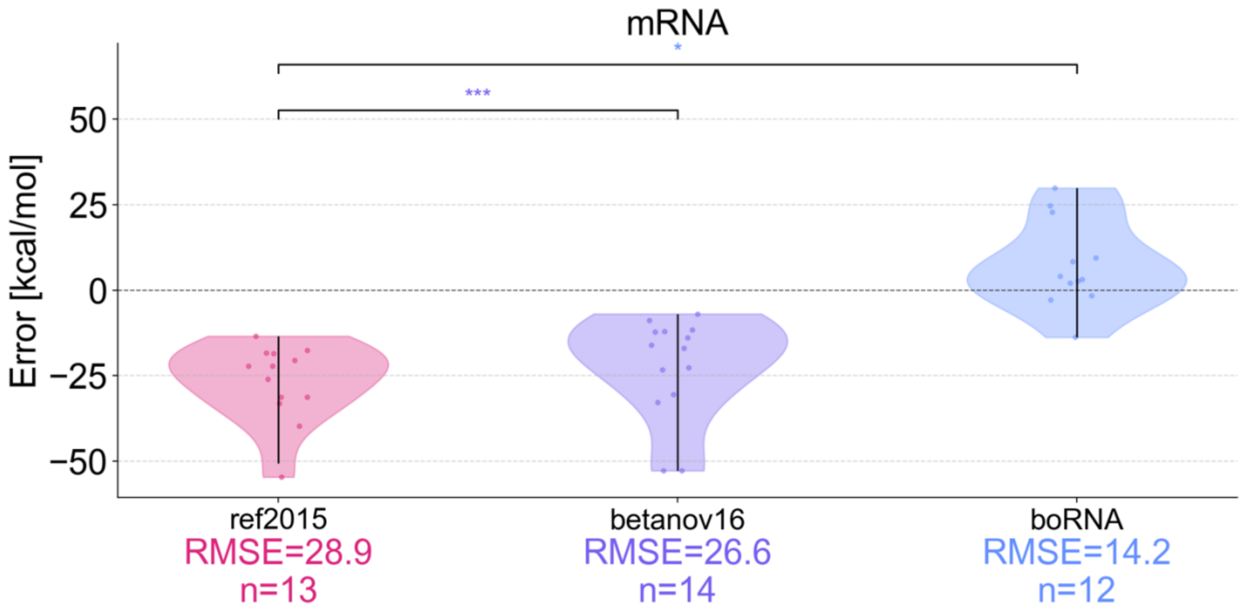


**Supplementary Figure 5.** mRNA Prediction Error.

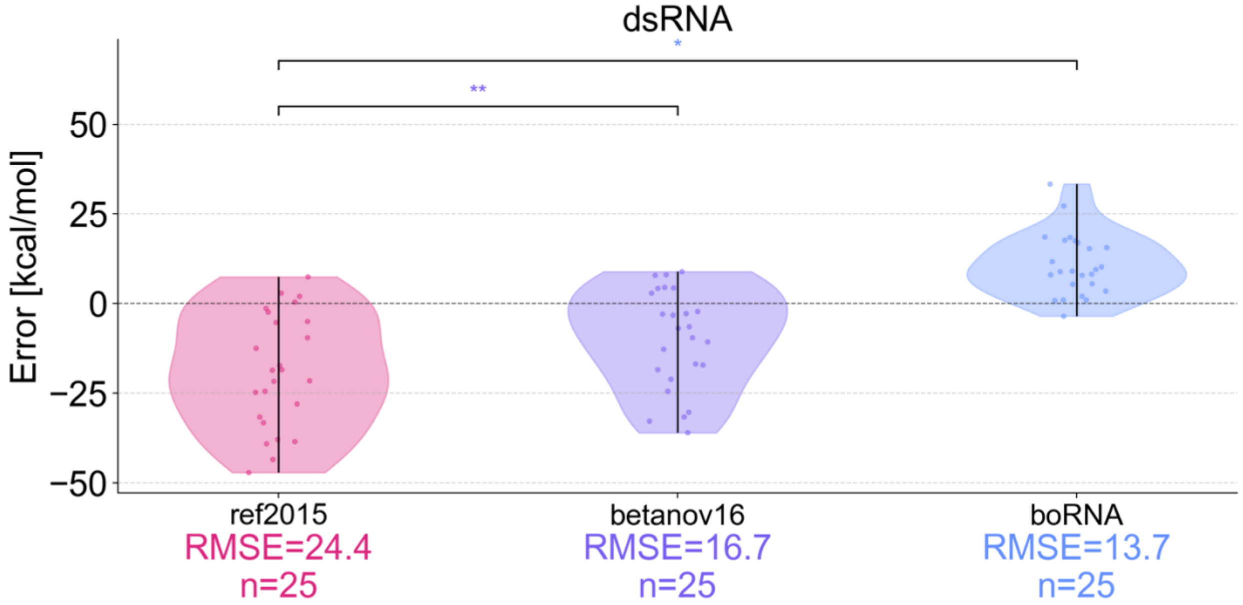


**Supplementary Figure 6.** dsRNA Prediction Error.

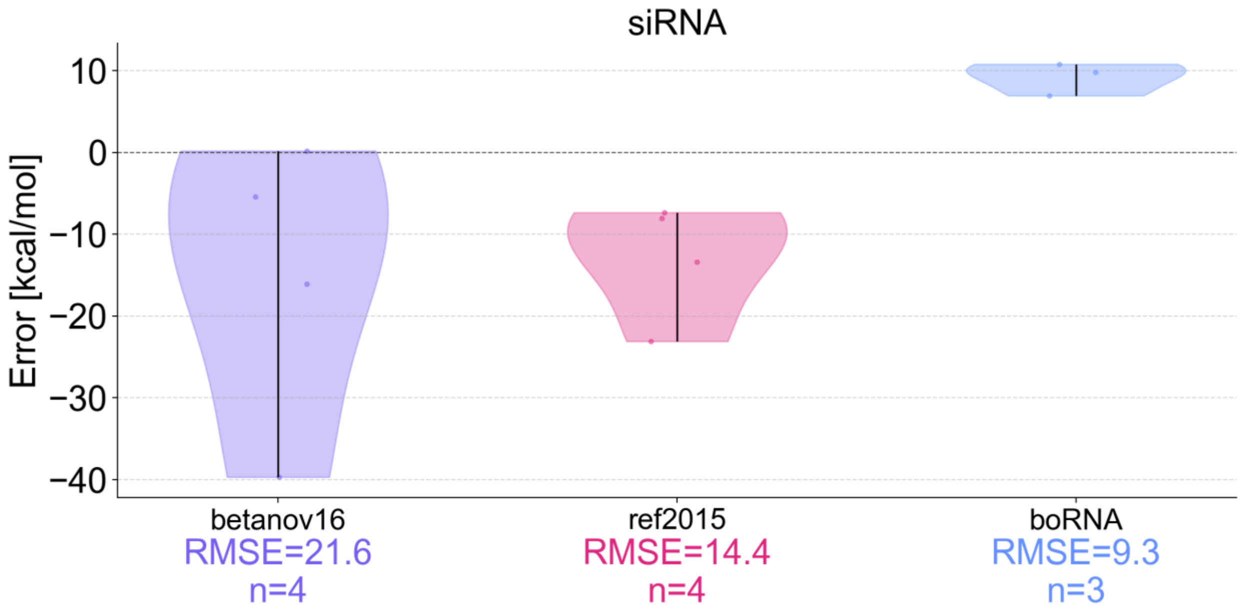


**Supplementary Figure 7.** siRNA Prediction Error.
